# Small-molecule screen in *C. elegans* identifies benzenesulfonamides as inhibitors of microsporidia spores

**DOI:** 10.1101/2025.02.25.640200

**Authors:** Qingyuan Huang, Haiyi Jiang, Junhong Wei, Yabin Dou, Guoqing Pan, Jie Chen, Aaron W. Reinke

## Abstract

Microsporidia, a large group of fungal-related intracellular parasites, infect several economically significant animals, leading to substantial economic losses. As currently available anti-microsporidia therapies are either ineffective or come with numerous adverse effects, there is a need for alternative microsporidia inhibitors. Here we screen a subset of the ChemBridge DIVERset library, comprising 2500 diverse compounds, using *Caenorhabditis elegans* infected with its natural microsporidian parasite, *Nematocida parisii*. By testing these compounds at 60 μM in 96-well assay plates, we identified 26 hits that restored the ability of *C. elegans* to produce progeny in the presence of *N. parisii*. We confirmed that out of 20 tested compounds, 18 ChemBridge compounds effectively inhibit *N. parisii* infection in *C. elegans*. Of these 18, 10 were benzenesulfonamide derivatives which inhibit microsporidia infection by inactivating spores. We screened an additional 475-compound benzenesulfonamide library, successfully identifying three compounds that are effective at a lower concentration than the initial hits. We further show that one benzenesulfonamide compound displays inhibitory activity against several species of microsporidia, inhibiting infection of species belonging to the *Nematocida*, *Enterocytozoon*, and *Encephalitozoon* genera. Together our results suggest that benzenesulfonamides are a potential scaffold for the development of microsporidia antiseptics.

## Introduction

Microsporidia are obligate intracellular pathogens that infect most kinds of animal groups^1–3^. Microsporidia were previously classified as protozoa, but they are now regarded as fungi^4^. Over 1700 species of microsporidia have been identified and they are widespread in the natural environment^5,6^. Microsporidian infections in host organisms are usually deleterious and can cause mortality^7,8^. A variety of economically significant animals, such as farmed penaeid shrimp, honey bees, and silkworms are at risk from these parasites^9,10^. Microsporidiosis outbreaks, especially in hatcheries, may collapse industries^11^. Additionally, 17 species of microsporidia have been reported to infect humans^12^. Patients with compromised immune systems, such as HIV-positive individuals, organ transplant recipients, or cancer patients, are more likely to be infected by this emerging pathogen^12^.

Anti-microsporidia therapies currently available are either ineffective or associated with several adverse effects^13^. Albendazole, a benzimidazole derivative, is one of the most common treatments for microsporidiosis. Other benzimidazole derivatives, such as MMV1782387, carbendazim, fenbendazole, and oxfendazole also show anti-microsporidia activity^14,15^. Benzimidazole analogs are effective in controlling microsporidiosis during the proliferative phase, however, they were not observed to enhance parasite clearance^15^. Benzimidazole analogs are effective against the intracellular stages of microsporidia, but ineffective on spores^16^. Albendazole is also less effective at inhibiting the human-infecting microsporidia *Encephalitozoon bieneusi* and *Vittaforma corneae*^17,18^. Albendazole is thought to bind to microsporidian beta tubulin, inhibit tubulin polymerization, and these microsporidian species contain variants that have been shown to cause albendazole resistance in other organisms^14^. The other most commonly used inhibitor of microsporidia is fumagillin, which binds specifically and covalently to methionine aminopeptidase type 2 (MetAP2)^19–21^. Fumagillin is not approved for human and agricultural use due to concerns regarding host toxicity^14,22^. Other inhibitors such as orlistat and dexrazoxane have been explored to a limited extent^23,24^. There remains a need to identify other inhibitors to treat microsporidia infections in both animals and agriculturally relevant species^15^.

Microsporidia can only grow inside of host cells, and it has been challenging to screen for inhibitors of this parasite. The model organism *Caenorhabditis elegans* has proven to be an effective model organism for drug discovery^25^. Several species of microsporidia infect this nematode, and studies using *C. elegans* have contributed to our understanding of microsporidian infection mechanisms and host immunity^26^. The most common species found infecting *C. elegans* is *Nematocida parisii*, which infects intestinal cells and leads to a decrease in size, reproductive capacity, and life expectancy^27–30^. *N. parisii* infection begins when spores are ingested and then use their unique infectious apparatus called the polar tube to deposit sporoplasms inside of intestinal cells. The sporoplasms then grow into meronts which differentiate into spores which are then defecated from the worms^26,31^. We recently developed a high-efficiency *C. elegans*-*N. parisii* drug screening system for the discovery and characterization of novel inhibitors of microsporidia^15,24^. Infection with *N. parisii* reduces the ability of *C. elegans* to produce progeny, allowing for a quantifiable phenotype that can be reversed with microsporidia inhibitors.

To identify novel microsporidia inhibitors, we screened 2500 compounds from the ChemBridge DIVERSet library. We identified and validated 18 compounds which could reduce the levels of *N. parisii* infection in *C. elegans*. Most of the inhibitors we identified were halide-containing benzenesulfonamides which inhibited microsporidia infection by inactivating spores. We show that one of these benzenesulfonamides exhibits broad-spectrum inhibitory effects on microsporidia, affecting species within the *Nematocida*, *Enterocytozoon*, and *Encephalitozoon* clades. We performed a targeted screen of other benzenesulfonamides, identifying several with increased affinity. Overall, this study identifies benzenesulfonamides as broad-spectrum inhibitors of microsporidian spores.

## Results

### Screen of ChemBridge DIVERset library identifies microsporidia inhibitors

To identify novel microsporidia inhibitors we screened 2500 small-molecule compounds from the ChemBridge DIVERset library collection^32^. Using our previously described 96-well infection assay, compounds were incubated with *N. parisii* spores. *C. elegans* at the earliest larval (L1) stage were added one hour later and then cultured in liquid at 21 °C for five days^15,24^. As a result of microsporidia infection, *C. elegans* produce fewer embryos. Compounds which inhibit microsporidia can restore the ability of the host animals to produce progeny. In order to quantify offspring production after incubation with spores and each compound, each well was stained with rose bengal and scanned using a flatbed scanner, followed by automated quantification of progeny number^33^. Each compound was screened in triplicate at a concentration of 60 µm (Data S1). Compared to uninfected controls, 26 compounds increased progeny production by 40% in infected worms (Fig. 1A). These compounds increased the ability to make progeny 2.4-17-fold compared to infected controls (Fig. S1 and Data S1).

**Fig. 1.**
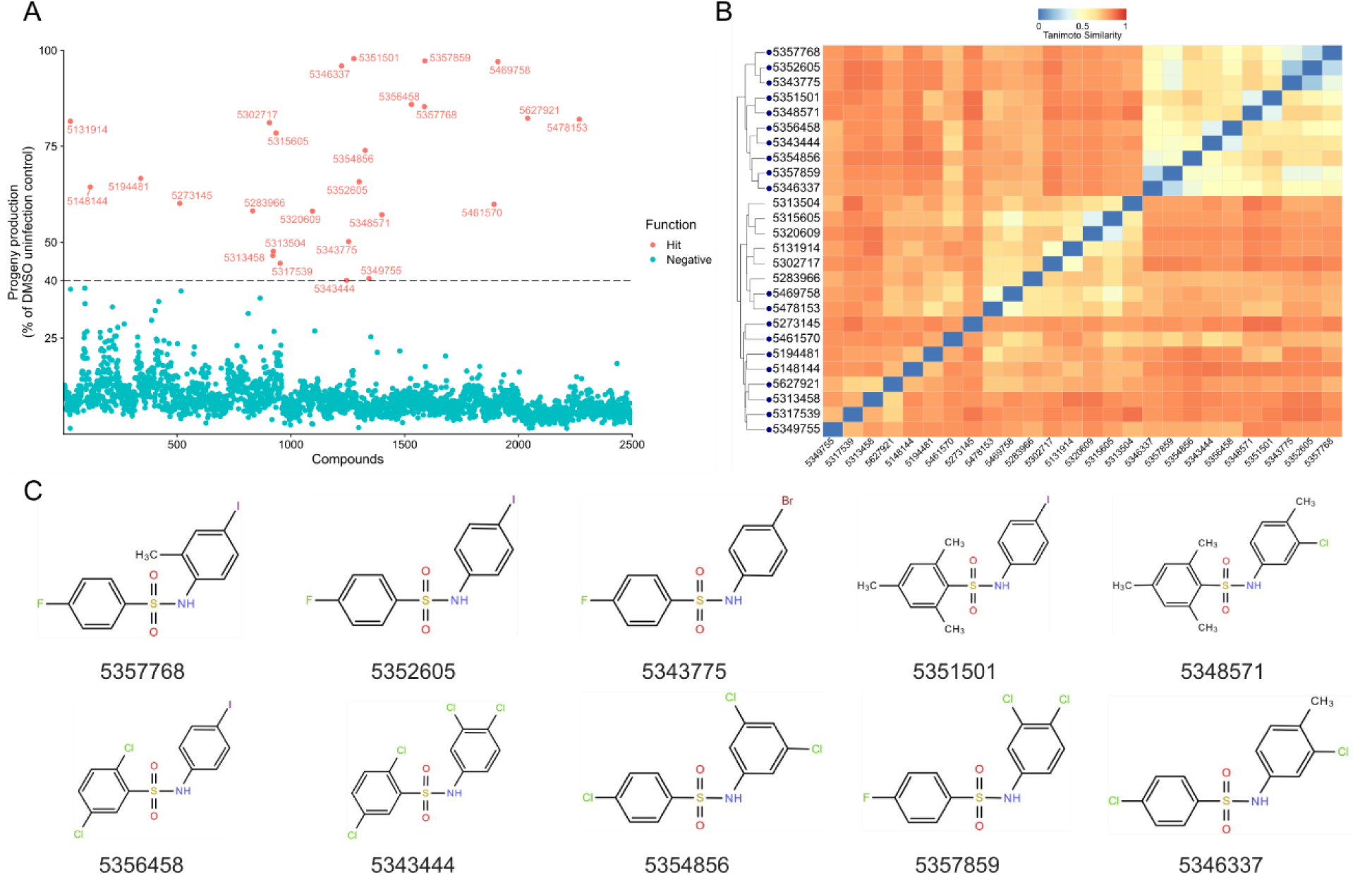
High-throughput screen of 2500 ChemBridge compounds identifies benzenesulfonamides which inhibit *N. parisii* infection of *C. elegans*. **A** Compounds at a final concentration of 60 µM were incubated with *N. parisii*. L1 stage *C. elegans* were added one hour later, cultured for five days, and progeny numbers quantified. Points represent mean progeny production of a compound as a percentage of DMSO uninfected controls. Compounds with an activity less than 40% are colored blue, compounds with an activity of at least 40% are colored red, and compound-IDs are shown. Three independent biological replicates were performed for each compound. **B** Clustered heat map of structural similarity of the 26 compounds with at least 40% activity. The scale indicates compound similarity with 0 (blue) being most similar and 1 (red) being least similar. Compounds tested in subsequent experiments are indicated by the dark blue dots. **C** Structures of benzenesulfonamides with inhibitory activity against *N. parisii*.

Hierarchical clustering analysis was used to determine the structural similarity of the identified microsporidia inhibitors (Fig. 1B). This analysis revealed a large cluster of ten compounds which are all benzenesulfonamide derivatives. All of these compounds contained both nitrogen- and sulfur-linked benzene rings. These 10 compounds also all contained at least 1 halide connected to a benzene ring (Fig. 1C). We also observed a second large cluster of compounds that contained either an aromatic imine or a pyrazole ring (Fig. S2). The other 8 compounds were mostly dissimilar from each other.

### Validation of compounds which inhibit *N. parisii* infection of *C. elegans*

To validate inhibitors identified in our screen, we selected 20 out of the 26 hits and quantified if the compounds could inhibit the infection of *N. parisii*. In a similar manner to our initial screen*, N. parisii* spores were cultured in the presence of compounds for one hour in 24-well plates, then L1 stage worms were added and cultured in liquid at 21 °C for four days. Afterwards, worms were fixed and stained with direct yellow 96 (DY96) which binds to both microsporidia spores and to *C. elegans* embryos^34^. In the presence of *N. parisii* spores, 18 of the 20 compounds, as well as the known microsporidia inhibitor dexrazoxane, significantly increased the proportion of adult worms containing embryos (which we refer to as gravid) (Fig. 2A)^24^. To determine the relationship between embryo numbers per worm from our validation experiment and progeny production from our screen, we performed a linear correlation which shows a moderate correlation with an R^2^ of 0.3044 (Fig. S3). Among the newly identified compounds, 16 of them significantly reduce the number of animals displaying newly formed *N. parisii* spores (Fig. 2B). Together our data shows that our high-throughput screen can identify inhibitors with high confidence. Additionally, all ten of the benzenesulfonamide inhibitors we identified significantly restore progeny production in infected *C. elegans* and 9 of them significantly inhibit *N. parisii* infection.

**Fig. 2.**
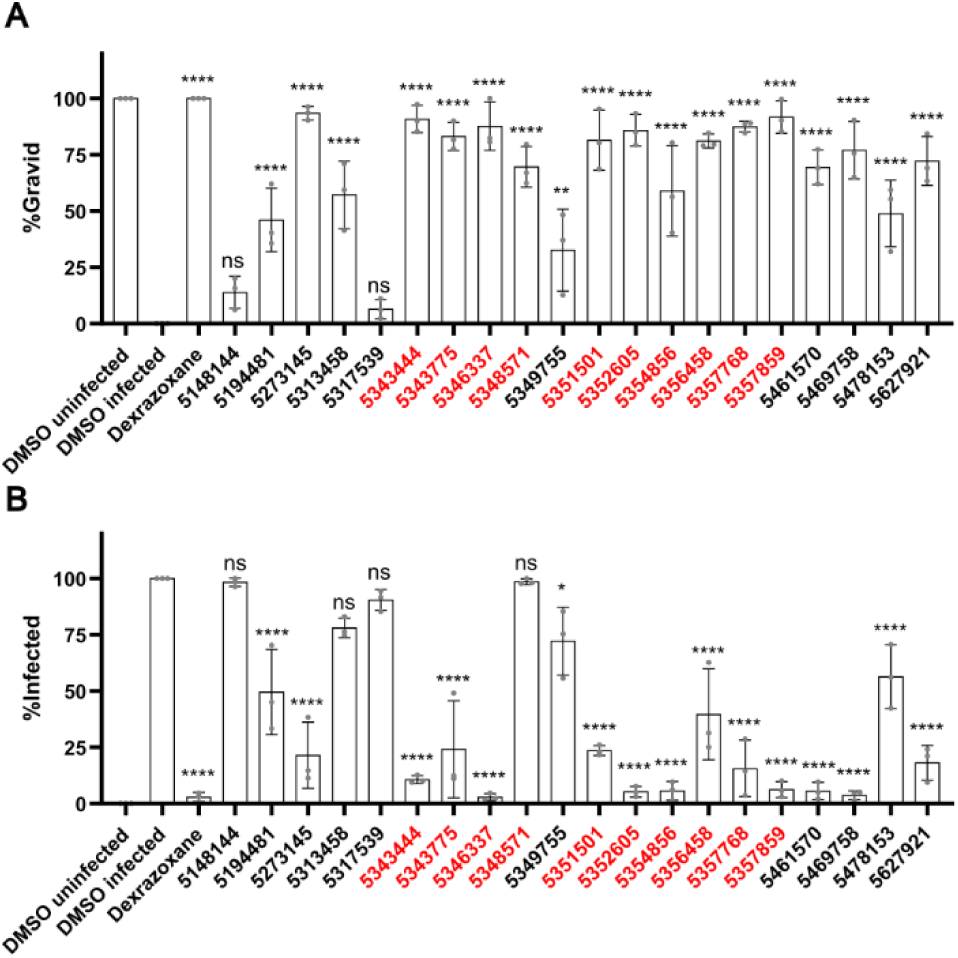
Validation that the identified ChemBridge compounds inhibit *N. parisii*. **A, B** *N. parisii* spores were incubated with compounds at a final concentration of 100 μM. One hour later, L1 stage worms were added and cultured for 4 days. Worms were then fixed and stained with DY96 to detect microsporidian spores and nematode embryos. **A** The percentage of animals which contain embryos. **B** The percentage of animals with newly formed spores. Benzenesulfonamide compounds are colored red. n = 3 biological replicates, N = ≥ 100 worms counted per biological replicate. The P values were determined by one-way ANOVA with post hoc test, with all comparisons to DMSO infected. Means ± SD (horizontal bars) are shown. (*p <0.05, **p < 0.01, ****p < 0.0001, ns means not significant).

Sulfonamides are a class of compounds that have been used as drugs in humans and animals to treat infectious disease^35^. We tested five medicinal sulfonamides for their activity against *N. parisii*. We did not observe any significant effect of these compounds in restoring embryo production of infected *C. elegans* or reducing infection levels (Fig. S4).

### Characterization of the mechanisms by which compounds inhibit *N. parisii*

Inhibitors of microsporidia could function by blocking infection or the ability of the intracellular stages of the parasite to proliferate inside of host cells. To determine if these compounds reduced microsporidia proliferation, we performed pulse-chase experiments. We infected the worms with spores, then washed them three hours later to remove non-ingested spores. Next, the infected worms were incubated with each of the 20 ChemBridge compounds, or dexrazoxane which has been previously shown to limit *N. parisii* growth^24^. One of the ChemBridge compounds, 5357859, increased the proportion of gravid worms, though to a much lower extent than dexrazoxane (Fig. 3A). None of the other 19 ChemBridge compounds reduced the percentage of worms displaying newly generated spores (Fig. 3B). To examine if ChemBridge compounds displayed inhibition of the meront stage of microsporidia growth, we stained meronts using Fluorescence *in situ* hybridization (FISH) at two days (before spore formation) or four days (after spore formation). Of the ChemBridge compounds, only 5357859 significantly reduced the amount of meronts in the worms, and to a much lesser extent than dexrazoxane (Fig. 3C and D). Similar to dexrazoxane, 5357859 did not decrease the rate of infection among animals, demonstrating that this compound does not promote pathogen clearance (Fig. 3E).

**Fig. 3.**
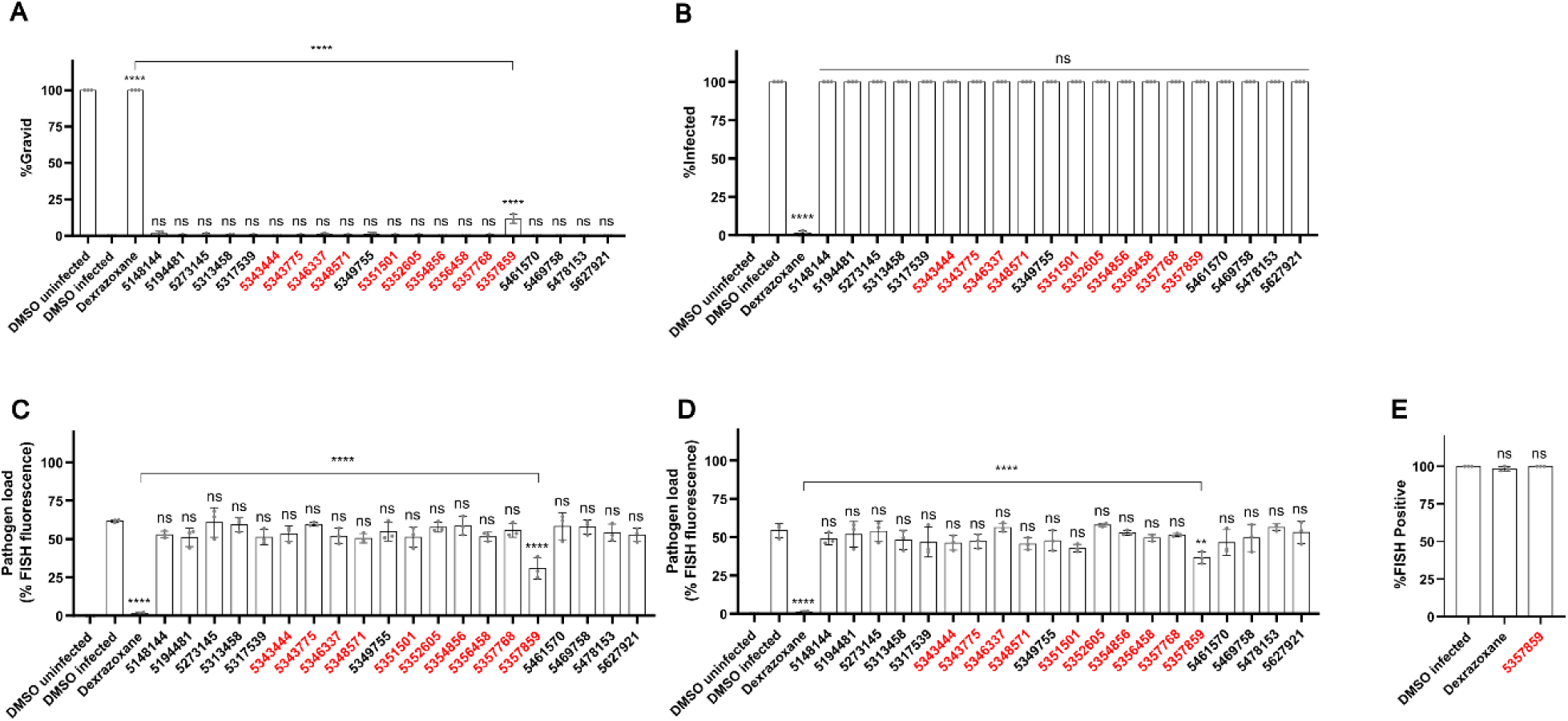
Tested ChemBridge compounds do not inhibit microsporidia proliferation except for 5357859. **A-E** L1 stage worms were incubated for 3 hours in the presence of *N. parisii* spores and then washed to remove undigested spores. Infected worms were incubated with compounds for 2 (D) or 4 (A-C, E) days, fixed, and stained with DY96 and a FISH probe specific to the *N. parisii* 18S rRNA. **A** The percentage of animals which contain embryos. **B** The percentage of animals which contain newly formed spores. **C, D** Quantification of the pathogen load at 2 (C) or 4 (D) days following infection. **E** The percentage of animals with FISH signal. Benzenesulfonamide compounds are colored red. n = 3 biological replicates, N = ≥ 100 worms (A, B, and E) or N=10 (C, D) counted per biological replicate. The P values were determined by one-way ANOVA with post hoc test, with all comparisons to DMSO infected, except for those indicated by brackets. Means ± SD (horizontal bars) are shown. (**p < 0.01, ****p < 0.0001, ns means not significant).

Dormant microsporidian spores can germinate (also called spore firing) within seconds of exposure to environmental stimuli^36^. In the intestinal lumen, many microsporidia, including *N. parisii*, germinate, depositing their sporoplasms inside of host cells, resulting in empty spores that no longer contain sporoplasms^37,38^. To determine whether any of the 18 ChemBridge compounds that had a significant effect on infected *C. elegans* progeny production also affect spore germination, we conducted spore firing assays^24^. Compounds were incubated with *N. parisii* spores for 24 hours, and then the spores were washed to remove the compounds. These treated spores were then used to infect L1 stage *C. elegans* for three hours. To visualize the sporoplasms and spores in the intestinal lumen, they were stained with both FISH and DY96. To determine the spore empty rate, we counted the number of empty spores divided by the total number of spores. ZPCK, a compound previously shown to inhibit spore germination^24^, and two ChemBridge compounds, 5313458 and 5343775, significantly reduced spore empty rates (Fig. 4A). Surprisingly, even though most of the compounds did not affect spore germination, the 18 ChemBridge compounds, as well as ZPCK, significantly decreased the amount of sporoplasms that invaded each worm (Fig. 4B). One possibility that would explain this incongruity is that the compounds could potentially decrease animal infection by inducing spore firing outside of the host. In order to determine whether any of the 18 ChemBridge compounds could induce spore firing *in vitro*, we incubated compounds with *N. parisii* spores for 24 hours, fixed and stained with DY96 and a *N. parisii* 18S rRNA FISH probe. None of the compounds significantly triggered spore germination *in vitro* (Fig. S5).

**Fig. 4.**
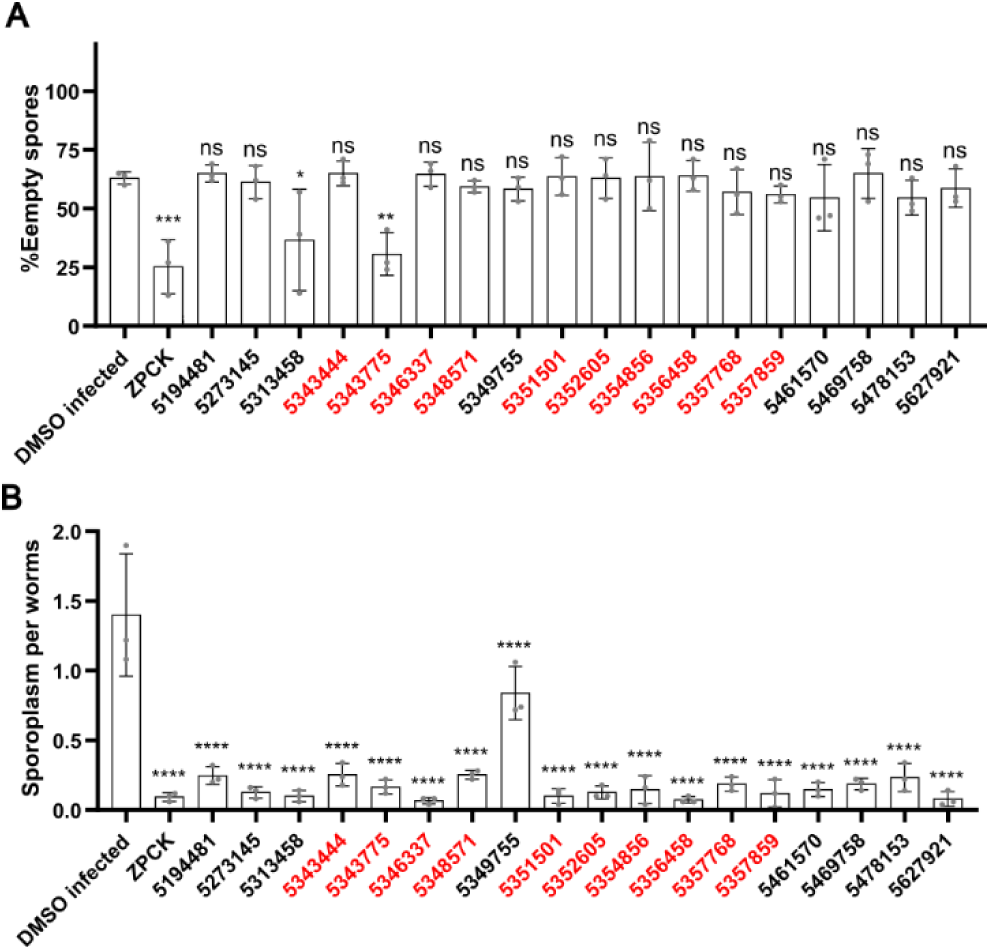
Identified ChemBridge inhibitors prevent *N. parisii* invasion. **A, B** After incubating *N. parisii* spores with compounds for 24 hours, spores were washed and added to L1 stage worms. After incubation for 3 hours, worms were fixed and the spores (DY96) and sporoplasms (FISH) were stained. **A** Percentage of empty spores in the intestinal lumen. **B** The mean number of sporoplasms per worm. Benzenesulfonamide compounds are colored red. (n = 3, N = ≥ 50 spores counted per biological replicate). The P values were determined by one-way ANOVA with post hoc test. Means ± SD (horizontal bars) are shown. (**p < 0.01, ***p < 0.001, ****p < 0.0001, ns means not significant).

### *N. parisii* spores are inactivated by benzenesulfonamide derivatives

One possibility that could explain how the identified compounds could block microsporidia invasion without altering the rate of empty spores, is if the compounds were lethal to the spores and facilitated the lysis of their sporoplasms by the intestinal contents of *C. elegans*. To test this hypothesis, we first performed mortality assays using the 18 Chembridge compounds and heat treatment as a control. After incubating the compounds with *N. parisii* spores for 24 hours, spores were stained with Calcofluor White M2R, which binds to the spores, and Sytox Green, which only stains the DNA of inviable spores. Heat treatment and 12 of the ChemBridge compounds, including all ten of the benzenesulfonamide derivatives, significantly increased spore mortality (Fig. 5A). We then tested if spores treated with either acetone or with several ChemBridge compounds could cause the spores to no longer have their sporoplasm. Spores were treated with acetone alone, compounds alone, or compounds followed by treatment with acetone. These spores were then either incubated with L1 stage *C. elegans* for three hours or analyzed *in vitro* for spore mortality. The treatment of spores with acetone or benzenesulfonamide derivatives results in a significant increase in spore mortality (Fig. 5B). In intestinal lumen, both 5343775 and 5357859 as well as their acetone co-treatment groups, show higher empty rates than the acetone treatment group (Fig. 5C). These results suggest that spores treated with the compound can lead to sporoplasm destruction in the intestinal lumen.

**Fig. 5.**
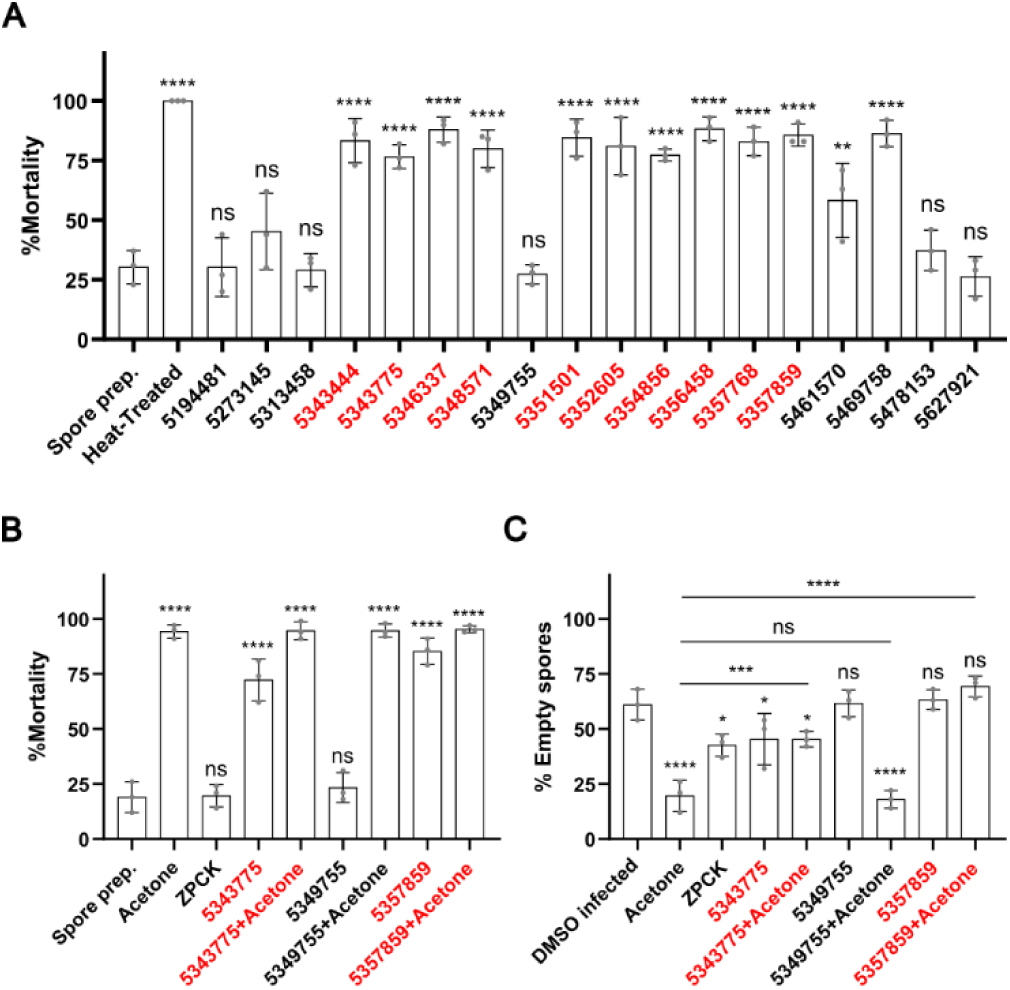
Benzenesulfonamides inactivate *N. parisii* spores. **A** *N. parisii* spores were treated with the indicated compounds for 24 hours, followed by Sytox Green and Calcofluor White M2R staining. Percentage of non-viable spores. **B**-**C** *N. parisii* spores were incubated with the indicated compounds, acetone, or both for 24 hours. Spores were then washed and stained with Sytox Green and Calcofluor White M2R (B) or mixed with L1 stage worms incubated for 3 hours, fixed, and stained with DY96 and a FISH probe (C). **B** Percentage of non-viable spores. **C** Percentage of empty spores in the intestinal lumen. N= ≥ 100 spores (A, B) or N = ≥ 50 worms (C) counted per biological replicate. The P values were determined by one-way ANOVA with post hoc test. Means ± SD (horizontal bars) are shown. (*p < 0.05, **p < 0.01, ***p < 0.001, ****p < 0.0001, ns means not significant).

### Multiple species of microsporidia are inhibited by benzenesulfonamides

In order to determine whether the compounds we identified are effective against other microsporidia species, we first tested them against *P. epiphaga*, which infects the epidermis and muscle of *C. elegans*. This species belongs to the *Enterocytozoonida* clade, which also contains the human-infecting species *V. cornea* and *E. bieneusi*^29,39,40^. FISH staining was used to determine whether the five selected ChemBridge inhibitors, including two benzenesulfonamides, could inhibit *P. epiphaga* infection in *C. elegans*. All of the tested compounds significantly inhibit *P. epiphaga* infection (Fig. 6A).

**Fig. 6.**
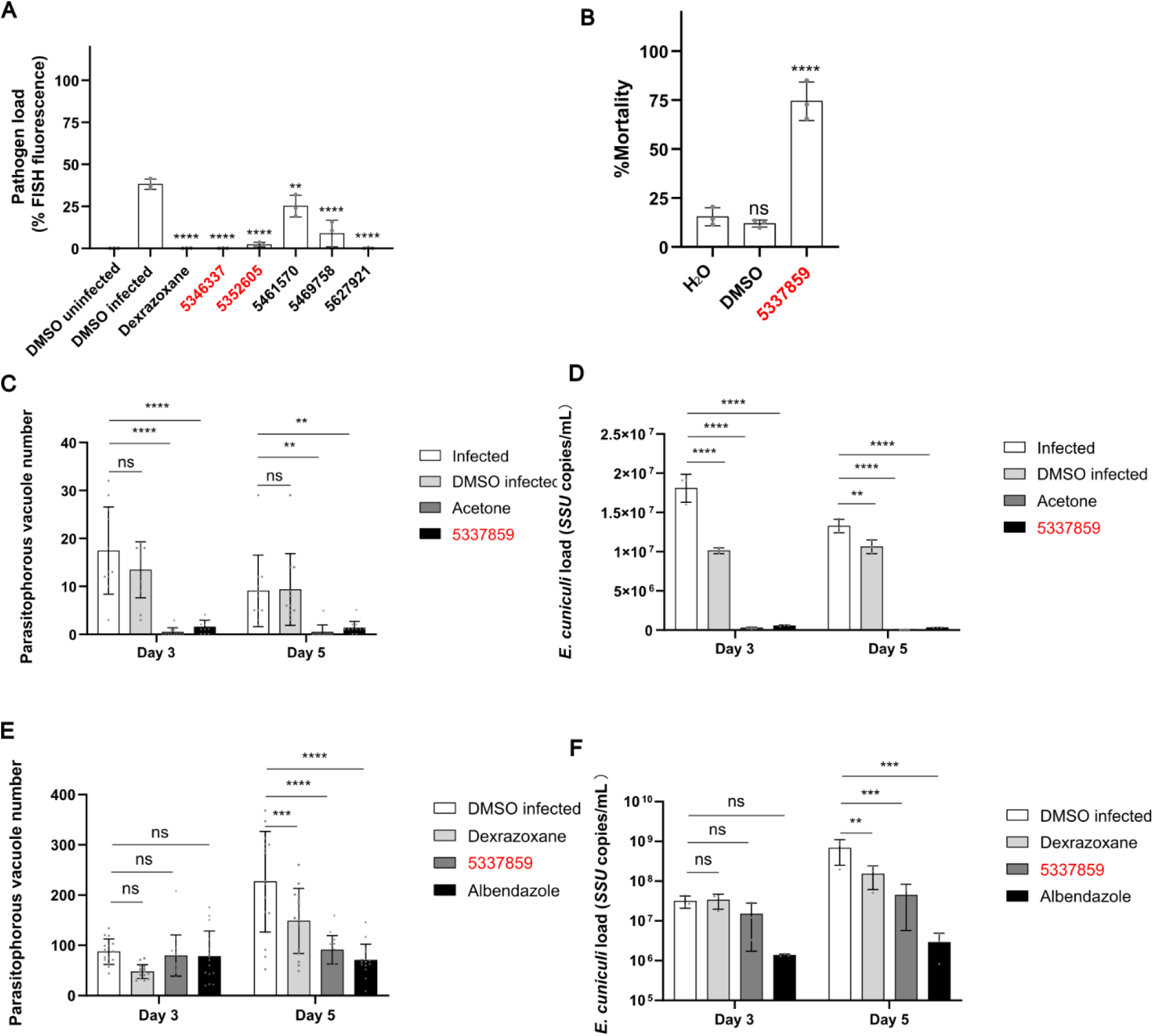
The benzenesulfonamide inhibitor 5357895 limits *P. epiphaga and E. cuniculi* infection. **A** *P. epiphaga* spores were incubated with compounds at a concentration of 200 μM for 1 hour and then L1 stage *C. elegans* were added and cultured for 4 days. Animals were then fixed and stained with DY96 and a *P. epiphaga* 18S rRNA FISH probe. Quantification of pathogen load. **B** *E. cuniculi* spores were incubated with 5337895, acetone, or 1% DMSO for 24 hours, followed by staining with Fluorescent Brightener 28, Sytox Green, and propidium iodine. Quantification of spore mortality. **C, D** *E. cuniculi* spores were incubated with 5337895, acetone, or 1% DMSO for 24 hours. Spores were then added to RK-13 cells and incubated for 3 days. **C** Cells were fixed, stained with Fluorescent Brightener 28 and parasitophorous vacuole number counted. **D** Quantitation of *E. cuniculi SSU* concentration. **E, F** RK-13 cells were incubated with *E. cuniculi* spores and either 5 μM dexrazoxane, 25 μM 5357859, or 200 nM albendazole for either 3 or 5 days. **E** Cells were fixed, stained with Fluorescent Brightener 28, and parasitophorous vacuole number counted. **F** Quantitation of *E. cuniculi SSU* concentration. Benzenesulfonamide compounds are colored red. n = 3 biological replicates, N = 10 animals (A), N= ≥ 100 spores (B), N >= 10 figure (C, E), or N = three well samples (D, F) qquantified per biological replicate. The P values were determined by one-way ANOVA (A-B) or two-way ANOVA(C-F) with post hoc test. Means ± SD (horizontal bars) are shown (*p <0.05, **p < 0.01, ****p < 0.0001, ns means not significant).

We then tested the ability of the benzenesulfonamide 5357859 to inhibit the microsporidian *Encephalitozoon cuniculi*, which can infect various hosts including farm and companion animals, as well as humans^41^. We first used mortality assays to determine that 5357859 effectively inactivates mature *E. cuniculi* spores *in vitro* (Fig. 6B). We treated spores with 5357859 and used them to infect RK-13 cells. We then measured infection levels using parasitophorous vacuole counting and real-time PCR analyses and found that these compounds significantly reduce the pathogen load after incubation for 3- or 5-days (Fig. 6C, D). To determine if treating *E. cuniculi* and cells at the same time could reduce infection, we incubated RK-13 cells with 5357859, albendazole, or dexrazoxane at the maximum safe dose we determined from cytotoxicity experiments (Fig. S6). After 3 days of incubation, we did not observe any significant decreases in parasitophorous vacuoles or Small Subunit (*SSU*) ribosomal RNA copies when treated with any of the compounds. After 5 days of incubation, we observed a significant decrease in the parasite by both measurements for dexrazoxane, albendazole, and 5357859 (Fig. 6E, F).

### Identification of higher affinity benzenesulfonamide analogs that inhibit *N. parisii* infection

To identify higher affinity benzenesulfonamide inhibitors, we screened a library of 475 ChemBridge benzenesulfonamide analogs. To identify benzenesulfonamide analogs with stronger inhibitory effects, we decreased the compounds concentration to 15 μM from the 60 μM that we used in the original screen. There were 7 benzenesulfonamides from the previous screen that served as negative controls, and none of them increased progeny production in infected worms by more than 18% (Data S1). We identified three compounds that increased progeny production in infected worms by more than 40% compared to the uninfected controls (Fig. 7A). Two of these compounds, 5351234 and 5354953, differ from 5357859 only by the position of the chlorines on the nitrogen-bound benzene. The other compound, 7819020, contains two fused benzene rings, with one of them connected to a bromine (Fig. 7B).

**Fig. 7.**
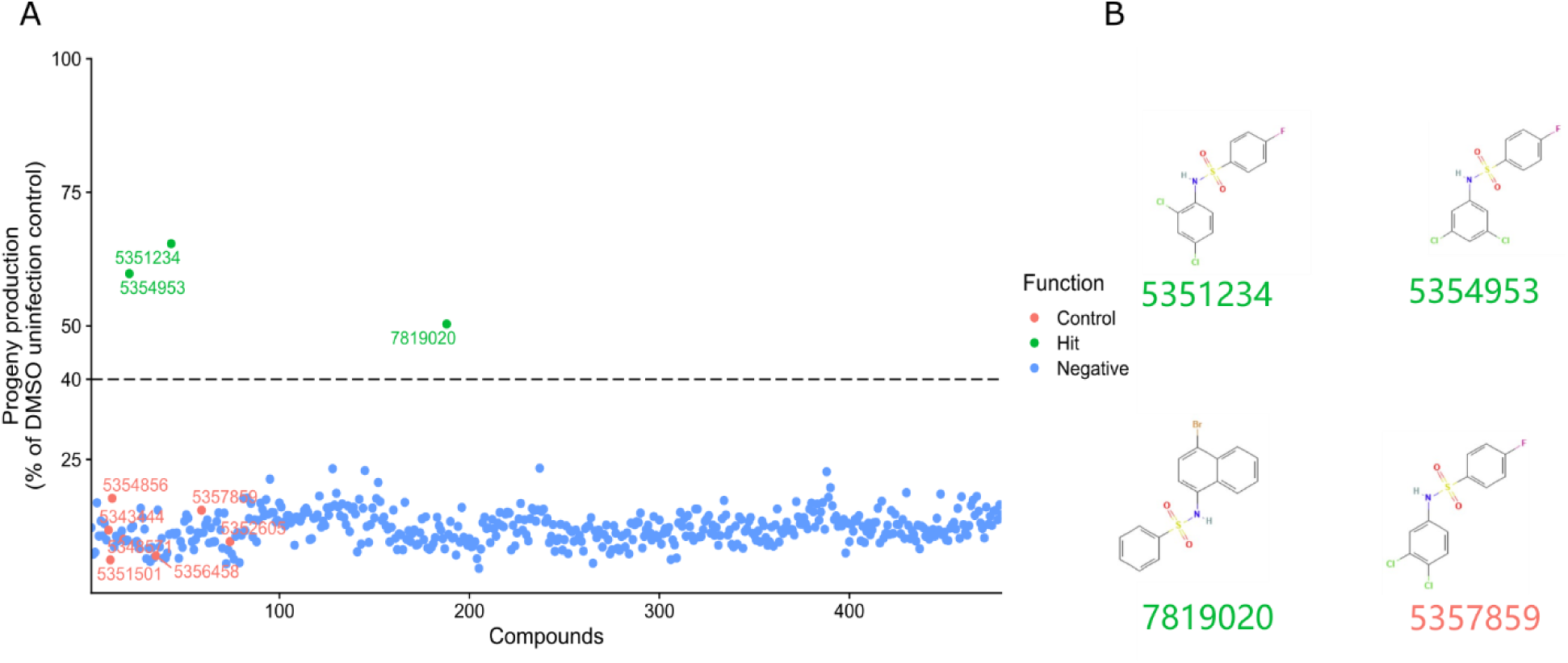
Screen of 475 ChemBridge benzenesulfonamide analogs identifies higher affinity *N. parisii* inhibitors. **A** Compounds at a final concentration of 15 µM were incubated with *N. parisii*. L1 stage *C. elegans* were added one hour later and then cultured for five days. The number of worms in each well were quantified and the progeny production of each compound as a percentage of the uninfected controls is shown. Three independent biological replicates were performed for each compound. The green sulfonamide-ID represents sulfonamides that exhibited an activity of at least 40%. The red sulfonamide-ID represents the activity of the previously identified inhibitor 5357859. **B** The chemical structures of 5357859 and the 3 ChemBridge benzenesulfonamides with higher affinity.

To validate and characterize the higher affinity benzenesulfonamide inhibitors, we treated both infected and uninfected worms with 5357859 and the three higher affinity benzenesulfonamides at a concentration of 15, 60 or 100 μM. To determine toxicity to *C. elegans*, we measured the proportion of uninfected animals that were gravid. None of the compounds significantly reduced the proportion of gravid animals at 15 μM (Fig 8A). However, 5351234 at 60 μM or higher, and 5354953 at 100 μM significantly reduced population gravidity (Fig 8E and 8I). We then determined the ability of these compounds to restore embryo production in animals infected with *N. parisii*. All of the compounds increased the percentage of gravid worms in the presence of *N. parisii* at 15, 60 or 100 μM (Fig 8B, 8F and 8J). At the lowest concentration 5357859 did not rescue gravidity to uninfected levels (Fig 8B). At the highest concentration 5351234 and 5354953 did not rescue gravidity (Fig 8J). We also evaluated whether these compounds could inhibit *N. parisii* infection, observing that all compounds mostly eliminated infection at the highest concentration, and that the three higher affinity benzenesulfonamides had higher activity than 5357859 at 15 μM (Fig 8C, 8G and 8K). Finally, we tested the ability of these compounds to cause spore mortality, observing that all compounds could and at 15 μM, 5357859 demonstrated a significantly reduced ability to inactivate spores than the other three (Fig 8D, 8H, and 8L).

**Fig. 8.**
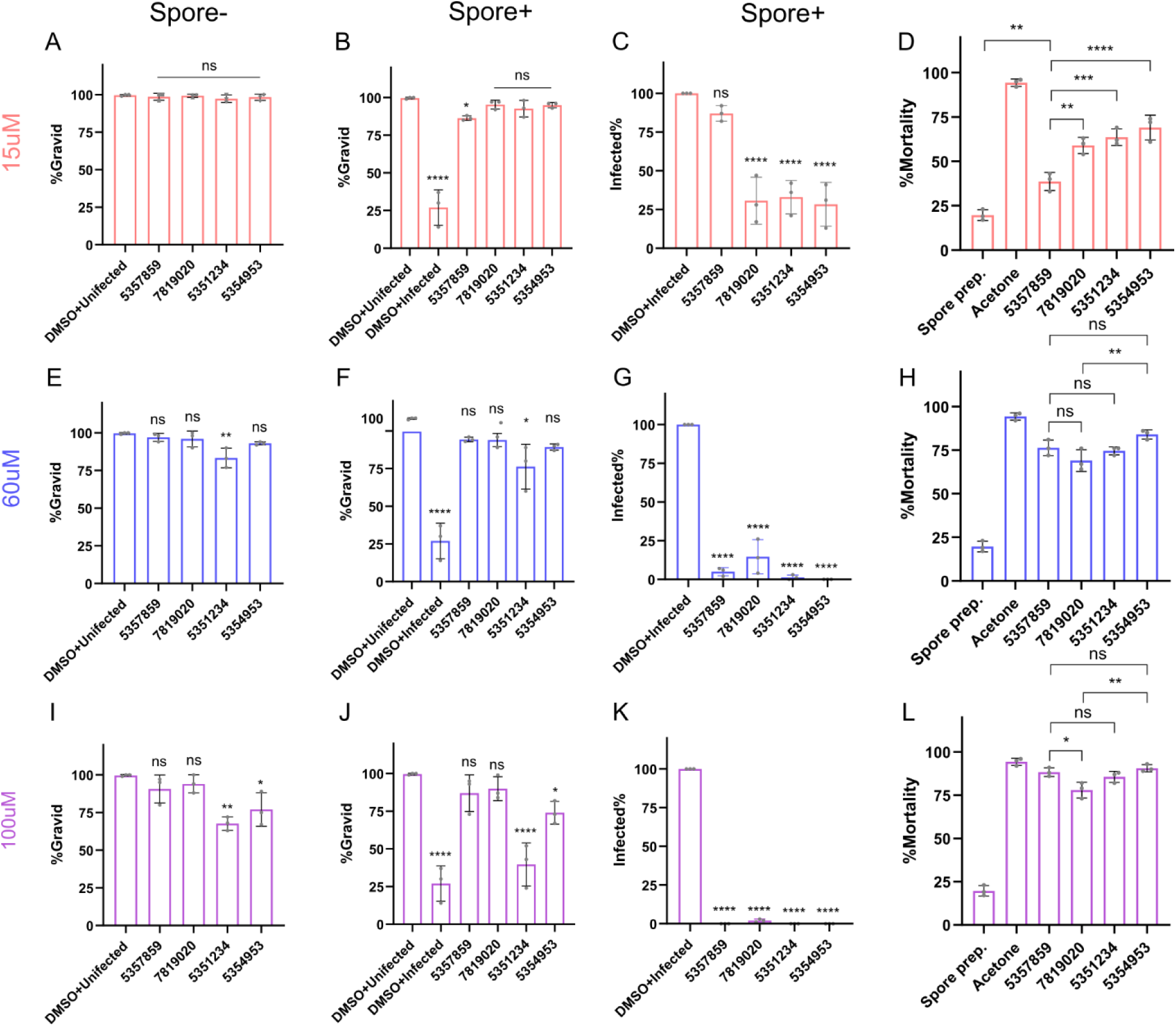
Higher affinity benzenesulfonamides inhibit *N. parisii* infection. **A-C, E-G and I-K** *N. parisii* spores were incubated with the indicated benzenesulfonamides at different concentrations for one hour. L1 stage *C. elegans* either without spore or mixed with the treated spores were cultured for four days. Worms were then fixed and stained with DY96. Effect of the indicated benzenesulfonamides at 15 µM (A-C), 60 µM (E-G) or 100 µM(I-K) on the percentage of worms with embryos in uninfected worms (A, E, and I), infected worms (B, F and J) and the percentage of worms with newly formed spores (C, F, and K). **D, H and L** *N. parisii* spores were incubated with 15 µM (D) 60 µM (H) and 100 µM (L) compound for 24 hours and stained with Sytox Green and Calcofluor White M2R. The percentage of spores that showed Sytox Green staining. n = 3, N = ≥ 100 worms (A-C, E-G, and I-K) or N= ≥ 100 spores (D, H and L) counted per biological replicate). The P values were determined by one-way ANOVA with post hoc test. Means ± SD (horizontal bars) are shown. (*p <0.05, **p < 0.01, ***p < 0.001, ****p < 0.0001, ns means not significant).

## Discussion

Through two rounds of high-throughput screening of a total of ∼3000 compounds, we identified 13 halide-substituted benzenesulfonamides, some of which can deactivate spores at low micromolar concentrations. We find that these compounds predominantly function by making spores non-viable and preventing invasion of host cells. We also find that these compounds have activity against several species of microsporidia. Although these compounds could potentially be used as antiseptics, at higher concentrations that cause nearly complete protection against infection, slight host toxicity is observed.

There has been effort to identify compounds that can be used as disinfectants or antiseptics to treat microsporidia spores^42^. Treatments such as hydrogen peroxide, ethanol, bleach, ozone, chlorine have been demonstrated to be effective against *Encephalitozoon* species^43–46^. However, some treatments, such as levulinic acid in combination with SDS, that are very effect against bacteria, do not inactivate microsporidia spores^47^. Several types of compounds have been shown to have activity on spores, but limited toxicity to hosts, such as porphyrins with *Nosema ceranae* or massetolides with *N. parisii*^48,49^. Most of the benzenesulfonamides we identified cause spore mortality, but do not induce or block germination. Almost all the benzenesulfonamides appear to only act on spores, but 5357859, also has slight infect on *N. parisii* proliferation. Although the mechanism of action is unknown, we speculate these compounds are causing membrane permeability that results in Sytox Green staining. The sulfonamide derivative OX11 containing halogen functional groups exhibited significant cell wall destruction and membrane rupture of *E. coli*, and the benzenesulfonamides we identified may be having a similar effect on microsporidia spores^50^.

Sulfonamides are an important class of medicinally active compounds, displaying a wide range of activities. Studies have indicated that sulfonamides act as antibacterial^51,52^, antifungal^51^, antiviral^53^, anti-tumour^54^, antimalarial^55^, anti-ulcer and anti-inflammatory agents^56^, as well as having anti-convulsant^57^ and anti-oxidant effects^58,59^. There are several case studies reporting treating microsporidia infections of humans with the sulfonamides sulfadiazine or sulfamethoxazole^60–62^. As these compounds were included as part of multi-drug regimens, their effectiveness in clinically treating microsporidia is unclear. We observe no activity of clinically used sulfonamides against microsporidia. Additionally, sulfonamides were reported to not have activity against *E. cuniculi*^63^.

In addition to benzenesulfonamides, we found several other types of compounds that were capable of inactivating *N. parisii*. A series of diaryl derivatives, such as (E)-2-{[(3-ethynylphenyl)imino]methyl}-4-nitrophenol), an analog of 5461570, was shown to specifically inhibit the interaction of NusB-NusE, decrease bacterial rRNA levels, and demonstrate antimicrobial activity against a variety of clinical methicillin-resistant *Staphylococcus aureus* strains without significant cytotoxicity against human cell lines^64^. A high-throughput screening process targeting Leishmania major revealed that 5469758 inhibits the parasite’s growth and vitality^65^.

In conclusion, we have identified benzenesulfonamides with the potential to be used as antiseptics. Being able to treat spores without host toxicity would have potential application against microsporidia that infect agriculturally important animals such as with honey bees, shrimps, and silkworms^9,10^. To further develop these compounds as antiseptics, it will be important to determine the mechanism by which these molecules inactivate spores and identify molecules with enhanced selectivity for microsporidia.

## Materials and Methods

### C. elegans maintenance

*Escherichia coli* OP50-1, the food source for *C. elegans*, was grown for 18 hours at 37 °C in lysogeny broth (LB) and then concentrated by centrifugation. The wild-type *C. elegans* strain N2 was cultivated as a mixed population, then L4 stage worms were picked onto nematode growth media (NGM) plates, which were seeded with 10x OP50-1 *E. coli* and kept at 21 °C for 4 days^66^. To synchronize the worms, M9 solution was used to wash them from the NGM plates, and sodium hypochlorite and sodium hydroxide were applied to bleach the worms. As soon as the embryos had been released from the gravid adults, they were washed and incubated at 21 °C for ∼18 hours to allow the embryos to hatch.

### N. parisii spore preparation

Stocks of *N. parisii* (ERTm1) spores were prepared as previously described^28,34^. On NGM plates, *C. elegans* N2 worms were infected with *N. parisii* spores. After incubating the infected worms for several days, they were harvested and frozen at −80 °C. Infected worms were mechanically disrupted using zirconia beads (2 mm diameter), followed by the removal of animal debris using a 5 μm filter (Millipore). Preparations of *N. parisii* spores were confirmed to be free of bacterial contamination and stored at −80 °C. The concentration of spores in the sample was determined by counting DY96-stained spores using a sperm counting slide (Cell-VU).

### Source of chemicals

ChemBridge DIVERSet screening compounds were acquired from SPARC Drug Discovery at the Hospital for Sick Children. These compounds were dissolved in 0.3 μl of DMSO at a concentration of 10 mM and arrayed into 96-well plates. For retesting, solid compounds were bought from ChemBridge. All compounds were stored at a temperature of −80 °C.

### Screening of microsporidia inhibitors in 96-well plates

Quantifying the ability of compounds to restore the ability of *C. elegans* to produce progeny in the presence of *N. parisii* was performed as previously described^15,24^. 96-well plates containing ChemBridge compounds in columns 2 to 11 were filled with 25 μl of K-medium (51 mM NaCl, 32 mM KCl, 3 mM CaCl2, 3 mM MgSO4, 3.25 μM cholesterol) containing 5x OP-50-1 and *N. parisii* spores, except for column 1, where spores were not added. 500 nl of DMSO was added to columns 1 and 12 for the DMSO uninfected and infected controls. We then added 25 μl of K-medium containing 24,000 spores/μl of *N. parisii* and plates were incubated at room temperature (∼21 °C) for 1 hour. Then, 25 μl of K-medium containing 100 bleach-synchronized L1 worms was added. After all the additions, each well contained 100 bleach-synchronized L1 worms, 1% DMSO, 12000 spores/μl *N. parisii* (except for uninfected controls) and 60 μM of compounds (except for infected and uninfected controls). Each 96-well plate was covered with a breathable adhesive porous film and placed inside humidity boxes wrapped in parafilm and incubated for five days at 21°C on a shaker with 180 revolutions per minute. Three independent biological replicates were performed for each compound, for a total of 96 plates being screened.

### Quantification of progeny production

After 5 days of incubation, a VIAFLO 96 Electronic pipette was used to add 10 μl of 0.3125 mg/mL Rose Bengal. Plates were wrapped in parafilm and incubated for 24 hours at 37°C, resulting in magenta staining of the worms. To each well, 240 μl of M9/0.1% Tween-20 was added and the plate was centrifuged for 1 minute at 2200 x g. Wells were washed again by removing 200 μl of supernatant from each well and adding 150 μl of M9/0.1% Tween-20. After mixing the worms in the plate by repeated pipetting, 25 μl of the liquid was transferred to another 96-well white clear bottom plates containing 300 μl M9/0.1%Tween-20. Worms in the plate were allowed to settle for 30 minutes and then the plates were scanned using an Epson Perfection V850 Pro flat-bed scanner with the following settings: positive film holder, 4800 DPI, and 24-bit color. The images were modified using GIMP version 2.10.18, with horizontal and vertical gridlines positioned such that each well is separated by a grid. HTML color codes #000000 and #FFC9AF were removed. Additionally, the images were modified by applying unsharp masking with the following parameters (radius = 10, effect = 10, threshold = 0.5). Yellow, blue, cyan, and green hue saturation were altered by adjusting lightness to 100 and saturation to −100. Red and magenta were adjusted to have a lightness of −100 and a saturation of 100. Using LZW compression, each well was exported as a single .tiff image. The number of worms was counted using WorMachine ^67^ running in MATLAB, with the pixel binarization threshold set to 30, the neighboring threshold set to 1, and the maximum object area was set to 0.003%.

### Structural similarity measurements

Atom Pair fingerprints (APfp) were used to calculate the structural similarity between the 26 compounds meeting our threshold for displaying anti-microsporidia activity^68,69^ The APfp of these compounds was determined using ChemMine tools^70^. Z-score values were used as the similarity measure and hierarchical clustering was performed^71^.

### Continuous infection assays

Compounds at a concentration of 100 μM, except for dexrazoxane which was at 60 μM, were incubated in 200 μl of K-medium containing 24,000 spores/μl of *N. parisii* for 1 hour. We then added 200 μl of K-medium containing 800 bleach-synchronized L1 worms to each well. Each well of 24-well assay plates contained a total volume of 400 μl including 800 L1 worms and 12,000 *N. parisii* spores/μl. Plates were then covered with adhesive porous film, placed inside a box, and the boxes were enclosed in parafilm. Plates were incubated for four days at 21°C using a shaker with 180 rpm. After incubation, the samples were washed twice with M9/0.1% Tween-20, fixed with acetone, stained with DY96, and analyzed by fluorescence microscopy.

### Pulse infection assays

To generate populations of infected worms, ∼8000 bleach-synchronized L1-stage worms, 30 million *N. parisii* spores, and 5 μl 10x OP50-1 were mixed and added to a 6-cm NGM plate. These plates were dried in a clean cabinet and then incubated for three hours at 21 °C. Worms were washed off of plates with M9/0.1% Tween-20, and then washed twice to remove undigested spores. Worms were added to 24-well plates. Each well of 24-well assay plates contained a total volume of 400 μl including 800 L1 worms with compounds present at a concentration of 100 μM, except for dexrazoxane which was at 60 μM. Three wells were tested for each biological replicate. Samples were incubated as described above, for either two or four days. Worms were then fixed in acetone and stained with DY96 and a FISH probe as described below.

### Spore firing assays

Spores at a concentration of 24,000 spores/μl, were incubated with compounds in 2% DMSO at a concentration of 200 μM except for ZPCK, which was at 120 μM, for 24 hours at 21 °C. The 24-well assay plate was prepared as described above after the spores were washed three times with K-medium. The assays concentrations were 12,000 spores/ μl, 100 μM compounds except ZPCK (60 μM), and 1% DMSO. After 3 hours of incubation, samples were fixed in acetone, stained with FISH and DY96, and observed with fluorescence microscopy.

### Mortality assays

*N. parisii* spores were incubated at a concentration of 24,000 spores/μl with the compounds at 200 μM in 2% DMSO for 24 hours at 21 °C. In the heat treatment group, the spores were heated at 100°C for 10 minutes. The spores were washed twice with 1 ml H_2_O, resuspended in 100 μl solution containing 2 μg/ml Calcofluor White M2R and 8 μM Sytox Green nucleic acid stain, and incubated at room temperature (∼21 °C for 10 minutes). A total of 2.5 μl of each mixture was then spotted onto slides, mixed with 2.5 μl of 2% agar, covered with a glass slide, and examined using fluorescence microscopy.

*E. cuniculi* spores were incubated at a concentration of 10,000 spores/μl with 200 μM compounds in 2% DMSO, 2% DMSO only, or acetone for 24 hours at 37 °C. The spores were washed twice with 1 ml PBS/0.1% Tween-20, resuspended in 100 μl solution of 0.5 μg/ml Fluorescent Brightener 28, 2.5 μM Sytox Green nucleic acid and 0.5 μM propidium iodine. The mixture was incubated for 10 minutes to facilitate staining, followed by examination using the Olympus BX53 microscope (Olympus, Tokyo, Japan).

### DY96 staining, fluorescence in situ hybridization (FISH), and fluorescence microscopy

After incubation, samples were washed twice with M9/0.1% Tween-20 to remove any remaining bacteria. Then the animals were fixed in 700 µl of acetone for 15 minutes and washed twice with PBS/0.1% Tween-20. DY96 solution (10 µg/ml DY96, 0.1% SDS in 1x PBS + 0.1% Tween-20) was added and samples were rotated for 30 minutes. The DY96 solution was then removed and EverBriteTM Mounting Medium with DAPI was added. For FISH staining, acetone-fixed samples were washed once with PBS/0.1% Tween-20 and then once with hybridization buffer (900 mM NaCl, 20 mM Tris HCl, 0.01% SDS) Samples were then incubated in the hybridization buffer containing 5 ng/µl microB *N. parisii* 18S rRNA FISH probe^72^ (ctctcggcactcctcctg) conjugated to Cal Fluor Red 610 (LGC Biosearch Technologies) for 24 hours at 46°C. Samples were washed once with 1 ml wash buffer (50 ml hybridization buffer + 5 mM EDTA). Samples were then stained with 20 µg/µl DY96. Zeiss Zen 2.3 was used to obtain images of the samples using a ZEISS Axio Imager 2 at magnifications ranging from 5x to 63x. Animals with any number of embryos were counted as gravid worms. An infected worm was defined as an animal that exhibits any newly formed spores.

### P. epiphaga infection assays

Stocks of *P. epiphaga* (JUm1396) spores were prepared as described above for *N. parisii*. The infection experiments with *P. epiphaga* were conducted using 24-well assay plates, with each well containing a final volume of 400 μl in K-medium, 800 L1 worms, and 60,000 *P. epiphaga* spores/μl. Except for dexrazoxane, which was used at 60 μM, all compounds were used at a concentration of 100 μM. The DMSO was used at a final concentration of 1%. Plates were incubated as described above for 4 days. The samples were then fixed in acetone and washed twice, stained with a CAL Fluor Red 610 FISH probe specific to *P. epiphaga* 18S rRNA (CTCTATACTGTGCGCACGG). Worms were imaged as described above and FISH fluorescence was quantified using the threshold and measure tools in ImageJ 2.9.0^73^.

### Cell culture and preparation of E. cuniculi spores

*E. cuniculi* spores (GB-M1) were propagated in RK-13 (rabbit kidney) cells as previously described^74^. The RK-13 cells were cultured in minimal essential medium supplemented with 10% fetal bovine serum and maintained at a constant temperature of 37 °C with 5% CO_2_. Following a cultivation period of 7 days, the RK-13 cells and spores were harvested. Cells were lysed by osmotic shock using water and pipetting to facilitate the release of spores. Cellular debris was removed using a 5 μm filter. Spore concentration was determined using a serial dilution method, followed by enumeration of the spores^75^.

### Infection of cells with pretreated E. cuniculi spores

100 μl containing 10 million *E. cuniculi* spores were incubated with 200 μM 5357859, 2% DMSO, or acetone for 24 hours at 37°C. The spores were then washed twice with 1 ml PBS/0.1% Tween-20. We then mixed 1×10^6^ spores with 1×10^5^ seeded RK-13 cells which were then added into each well of 12-well plate. The medium in the well was refreshed every 3 days. At 0, 3 or 5 days after incubation, the culture media was removed and washed twice with PBS and the cells were harvested for microscopy or qPCR detection.

### Continues infection with E. cuniculi spores

After seeding 5×10^4^ RK-13 cells in a 24-well plate, medium containing 5 μM dexrazoxane, 25 μM 5357859, or 200 nM albendazole was added into wells. After 8 hours, 2.5×10^5^ *E. cuniculi* spores were added to each well. The medium in the wells was refreshed every 3 days. After either 3 or 5 days after incubation with spores, the culture media was removed and the wells were washed three times with PBS, and the cells were harvested for microscopy or qPCR detection.

### Parasitophorous vacuoles counting

After infection of RK-13 cells with *E. cuniculi* spores either 3 or 5 days, the culture medium was discarded, and the cells were washed three times with PBS/0.1% Tween-20. Cells were then fixed in 4% paraformaldehyde for 15 minutes and washed 3 times with PBS/0.1% Tween-20 to remove excess fixative. 0.5 μg/ml Fluorescent Brightener 28 working solution was then added to the samples and incubated in the dark for 30 minutes. The cells were then washed three times with PBS/0.1% Tween-20. The number of parasitophorous vacuoles was then examined and counted under a AX10 fluorescence microscope (Germany).

### SYBR Green Real-time PCR

DNA was extracted from cells utilizing the ONEGA E.Z.N.A. Tissue DNA Kit. Absolute fluorescence quantification PCR was conducted in accordance with previously described methods (Forward primer ECUNF1: 5’-TCCTAGTAATAGCGGCTGACGAA-3’, Reverse primer ECUNR2: 5’-ACTCAGGACTCAGACCTTCCGA-3’) ^76^. The qPCR reactions were carried out using the SYBR Green qPCR Kit. All real-time PCR experiments were conducted in triplicate using the FQD-96A Fluorescence Quantitative Polymerase Chain Reaction Detection System (Bioer Technology, Hangzhou, China)

### Analyses of Cell Viability

Cell viability was assessed using the Cell Counting Kit-8 (CCK-8). A total of 10,000 cells per well, along with the compounds, were seeded into 96-well plates. At the tested time points, 10 μl of CCK-8 reagent (C0041, Beyotime, China) was added to every well as per the manufacturer’s instructions. The cells were then incubated for an additional hour at 37 °C with 5% CO_2_. Finally, the optical density was measured at 450 nm.

### Statistical analyses

The data were collected from three independent biological repeats and analyzed by GraphPad Prism 8. P-values were determined by ANOVA. Statistical significance was defined as p<0.05, *p<0.01, ***p<0.001, and ****p<0.0001.

## Supplementary materials

**Fig. S1.**
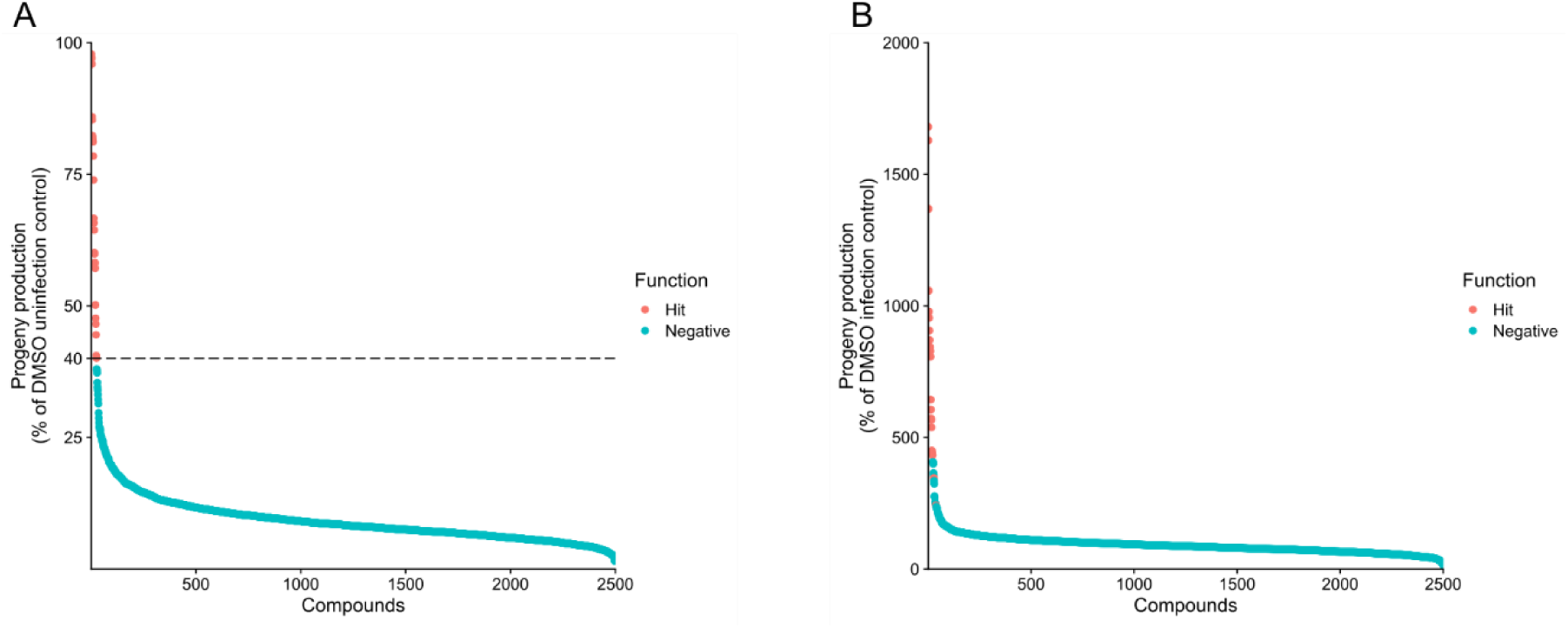
Ranked progeny production and fold increase for 2500 ChemBridge compounds tested against *N. parisii* infected *C. elegans.* **A, B** Data described in Figure 1 is presented in ranked order. Compounds with progeny production less than 40% are shown in blue and compounds with progeny production of at least 40% are shown in red. The data for each compound is displayed as the percentage of progeny production compared to uninfected controls (A) or as the percentage of progeny production compared to infected controls (B).

**Fig. S2.**
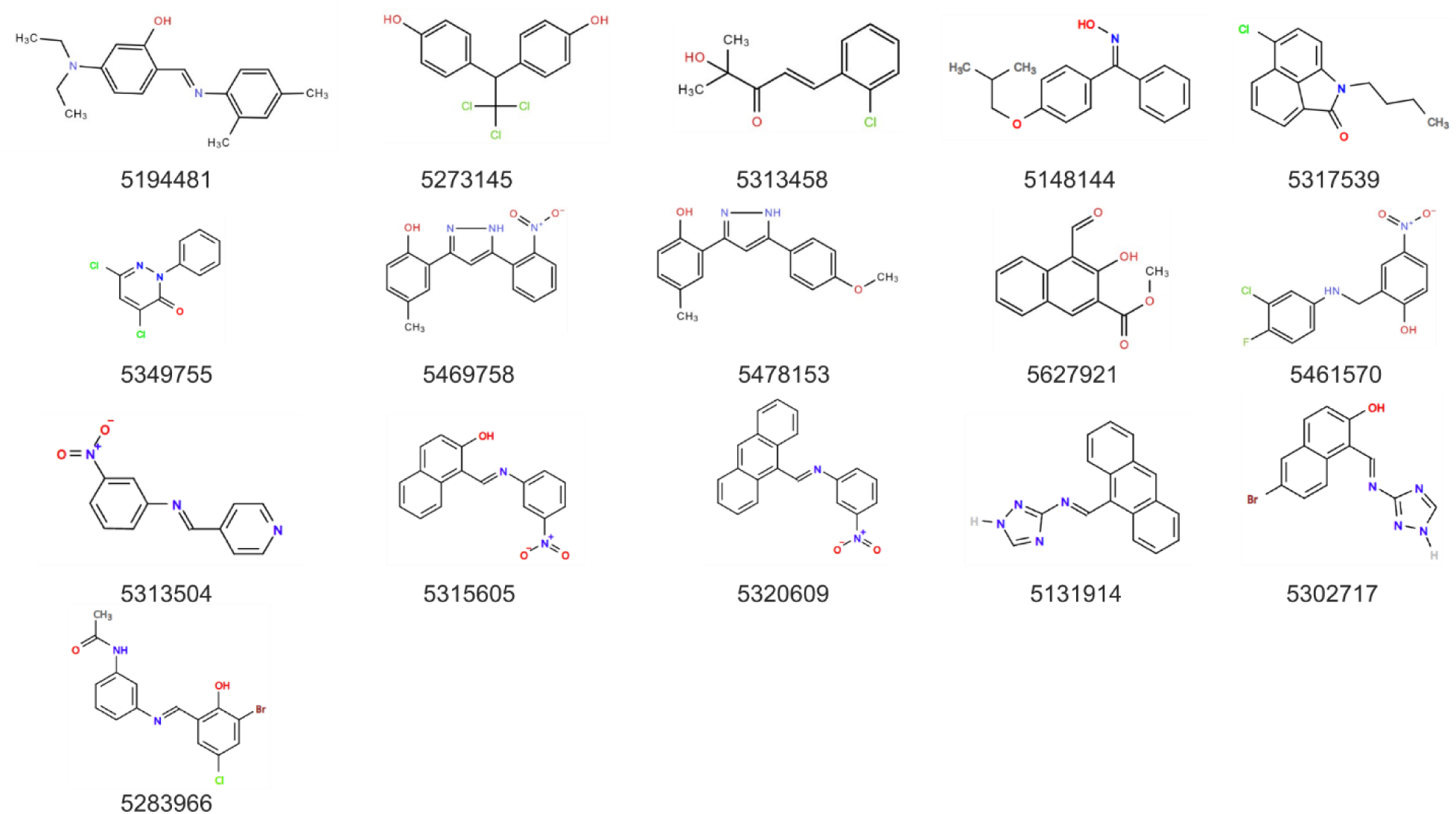
The chemical structures of the 16 non-benzenesulfonamide ChemBridge compounds with progeny production of at least 40%. The structure of each compound is shown with the label below the compound.

**Fig. S3.**
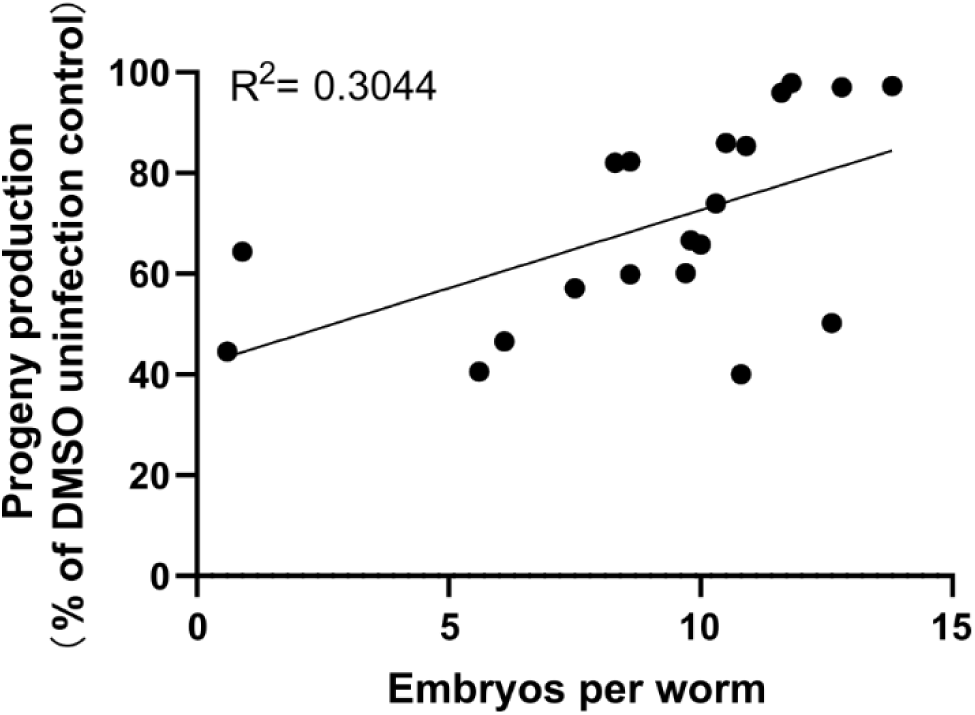
Progeny production from high-throughput screen is modestly correlated with number of embryos per worm. A linear correlation was performed between the number of embryos per worm for the 20 compounds tested in Figure 2A and the percentage progeny production from Figure 1A.

**Fig. S4.**
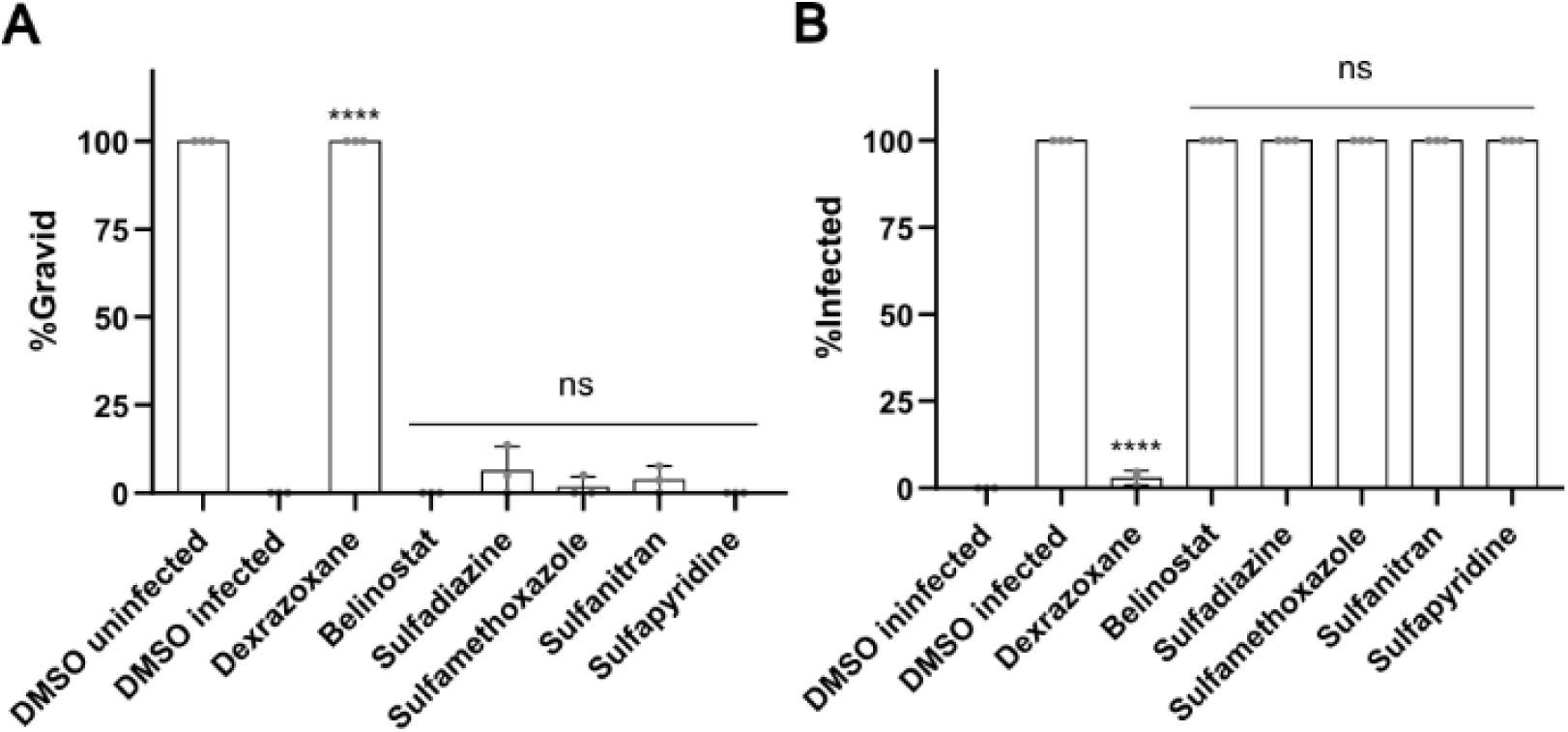
Medicinal sulfonamides do not inhibit *N. parisii*. **A, B** *N. parisii* spores were incubated with compounds at a concentration of 200 μM. One hour later, L1 stage *C. elegans* were added and cultured for four days. Following fixation, worms were stained with DY96. **A.** The percentage of animals which contain embryos. **B** The percentage of animals with newly formed spores. n = 3 biological replicates, N = ≥ 100 worms counted per biological replicate The P values were determined by one-way ANOVA with post hoc test. Means ± SD (horizontal bars) are shown. (****p < 0.0001, ns means not significant).

**Fig. S5.**
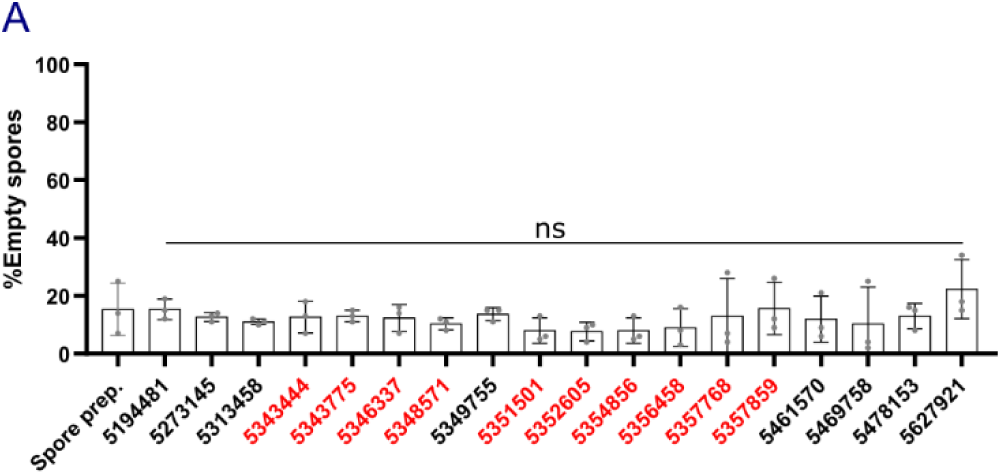
Identified ChemBridge compounds do not trigger sporing firing *in vitro*. **A** After incubating *N. parisii* spores with compounds for 24 hours, the compounds were washed away from the spores. Afterwards, spores were fixed and stained with FISH probe specific to the *N. parisii* 18S rRNA, DY96 and DAPI. Percentage of empty spores (n = 3, N = ≥ 100 spores counted per biological replicate). The P values were determined by one-way ANOVA with post hoc test. Means ± SD (horizontal bars) are shown. (ns means not significant).

**Fig. S6.**
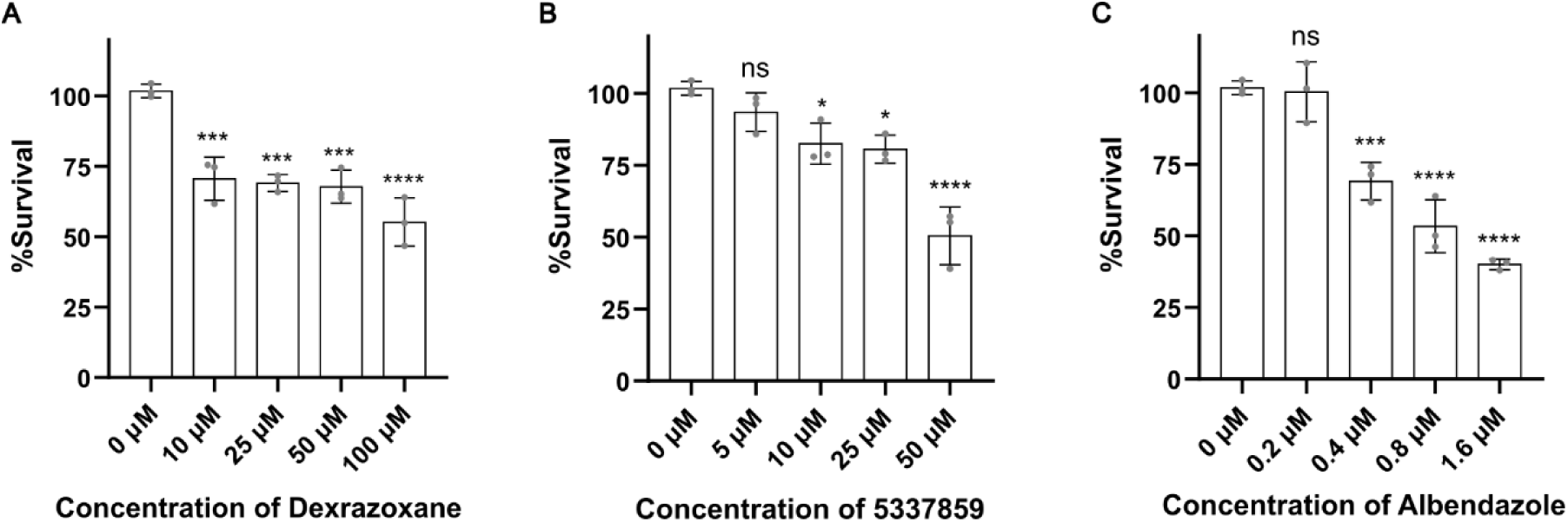
Determination of RK-13 cell viability when treated with microsporidia inhibitors. **A-C** The RK-13 survival rate was detected at different concentrations of either dexrazoxane (A), 5337895 (B), or albendazole (C). n= 3 biological replicates, N = 3 wells. The P values were determined by one-way ANOVA with post hoc test. Means ± SD (horizontal bars) are shown (*p <0.05, ***p < 0.001, ****p < 0.0001, ns means not significant).

